# Best Practices in Docking and Activity Prediction

**DOI:** 10.1101/039446

**Authors:** Manuel Rueda, Ruben Abagyan

## Abstract

During the last decade we witnessed how computational docking methods became a crucial tool in the search for new drug candidates. The ‘central dogma’ of small molecule docking is that compounds that dock correctly into the receptor are more likely to display biological activity than those that do not dock. This ‘dogma’, however, possesses multiple twists and turns that may not be obvious to novice dockers. The first premise is that the compounds must dock; this implies: (i) availability of data, (ii) realistic representation of the chemical entities in a form that can be understood by the computer and the software, and, (iii) exhaustive sampling of the protein-ligand conformational space. The second premise is that, after the sampling, all docking solutions must be ranked correctly with a score representing the physico-chemical foundations of binding. The third premise is that ‘correctness’ must be defined unambiguously, usually by comparison with ‘static’ experimental data (or lack thereof). Each of these premises involves some degree of simplification of reality, and overall loss in the accuracy of the docking predictions.

In this chapter we will revise our latest experiences in receptor-based docking when dealing with all three above-mentioned issues. First, we will explain the theoretical foundation of ICM docking, along with a brief explanation on how we measure performance. Second, we will contextualize ICM by showing its performance in single and multiple receptor conformation schemes with the Directory of Useful Decoys (DUD) and the Pocketome. Third, we will describe which strategies we are using to represent protein plasticity, like using multiple crystallographic structures or Monte Carlo (MC) and Normal Mode Analysis (NMA) sampling methods, emphasizing how to overcome the associated pitfalls (e.g., increased number of false positives). In the last section, we will describe ALiBERO, a new tool that is helping us to improve the discriminative power of X-ray structures and homology models in screening campaigns.

## Introduction

During the last decade we witnessed how computational docking methods became a crucial tool in the search for new drug candidates. The ‘central dogma’ of small molecule docking is that compounds that dock correctly into the receptor are more likely to display biological activity than those that do not dock. This ‘dogma’, however, possesses multiple twists and turns that may not be obvious to novice *dockers*. The first premise is that the compounds must dock; this implies: (i) availability of data, (ii) realistic representation of the chemical entities in a form that can be understood by the computer and the software, and, (iii) exhaustive sampling of the protein-ligand conformational space. The second premise is that, after the sampling, all docking solutions must be ranked correctly with a score representing the physico-chemical foundations of binding. The third premise is that ‘correctness’ must be defined unambiguously, usually by comparison with ‘static’ experimental data (or lack thereof). Each of these premises involves some degree of simplification of reality, and overall loss in the accuracy of the docking predictions.

## 1. Ligand Sampling with ICM

Docking with ICM (Internal Coordinate Mechanics)^3^ relies on exhaustive exploration of the degrees of freedom of the ligand within the boundaries of a “static” receptor. The receptor (or pocket) is described as a set of potential grid maps, representing van der Waals potentials for hydrogens and heavy atoms, electrostatics, hydrophobicity, and hydrogen bonding. Before the docking, the ligands are placed in the pocket in several random orientations that will serve as starting points for the MC sampling. During the sampling, ligand internal coordinates are optimized using biased probability Monte Carlo (BPMC)^4^ with an energy function that includes the ligand internal strain and a weighted sum of the grid map values in ligand atom centers. After the MC, a given number of binding poses (usually between 1-5) are evaluated with an all-atom ICM empiric ligand binding score derived from a multi-receptor screening benchmark by obtained as a compromise between approximated Gibbs free energy of binding and numerical errors. On average, a flexible-ligand ICM docking run takes ~ 30-60 seconds per ligand.

## 2. Measuring Docking Performance

The goal of small molecule docking is to predict the geometry of a ligand (a chemical compound) along with their interactions with a receptor (normally a protein). This objective can be subdivided into two overlapping categories: (i) ligand pose prediction, and, (ii) ligand screening. Pose prediction (a.k.a. geometry prediction or binding mode prediction) relies on the ability to reproduce the experimental coordinates of a ligand. If the coordinates of the complexes are available (i.e., coming from X-ray crystallography or nuclear magnetic resonance (NMR)), then they can be used retrospectively to check the predictions. Ligand screening, on the other hand, does not rely per se in geometry but rather in measuring the capacity of discrimination of true active molecules from inactives.

### 2.1 Pose prediction

Today, most of the sources of three-dimensional (3D) information for structure-based drug design (SBDD) are based on X-ray crystallography. Consequently, most docking tools are parameterized, tuned and benchmarked by their capacity to reproduce the 3D coordinates of *known* X-ray structures.

Probably the most common way of checking the correctness of docking results is by computing the root mean squared deviation (RMSD). The RMSD accounts for the average distance of the ligand atoms in the model with respect to the atoms in the reference structure after receptor superimposition. Normally, a ligand pose at ≤2 Å RMSD heavy-atom distance with respect to the crystallographic pose is accepted as “near-native” solution.^5^ Despite its wide acceptance, RMSD possesses shortcomings leading to misclassification of correct and incorrect poses.^5,6^ For instance, RMSD requires receptor superposition which is rather ambiguous when models represent distinct conformations or when the proteins are non-identical. Any receptor superimposition implies (i) judicious choice about the portion of the protein to be superimposed, (ii) atomic equivalence in the selection (note that symmetry in side chain atoms may be involved), and, (iii) down-weighting the contribution of flexible or poorly defined regions. In that regard, a robust method that iteratively optimizes the superimposition by assigning lower weights to most deviating fragments is preferred.^7^ Ligands are also affected by RMSD idiosyncrasies. For example, once the receptors have been superimposed, atomic equivalence must be established for ligand atoms, which is not a trivial task. On top of that, 2D to 3D ligand transformations can create conflicts in naming / atom type schemes even *within* the same docking platform. Even if the “chemistry” has not been distorted by the software technicalities, a RMSD still may not be indicative of how well the critical interactions are conserved. For instance, large RMSD values can come from a compound displaying symmetry elements totally flipped over,^6, 8^ or from flexible atoms not interacting with the protein, like the solvent-exposed parts of the ligand in the GPCR Dock 2008 assessment^9, 10^ (the solvent exposed phenoxy group of the adenosine A_2A_ receptor antagonist; PDB ID: 3EML).

As an alternative to the RMSD, *interaction-based* measures reflect molecular recognition by measuring key protein-ligand contacts. As a rule, interaction-based measures account for geometrical distances between atoms in the receptor and atoms in the ligand, and have the advantage of being invariant to receptor superimposition.^6, 11^ Unfortunately, despite their utility, we realized that even the simplest comparison (i.e., comparing the equivalent contacts at a given distance cutoff with respect to an X-ray structure) could become extremely challenging even with the right tools on hand. As with the RMSD, atomic equivalence must be established, but this time, to make things worse, neighboring side chain atoms may display symmetry leading easily to exponential growth of complexity. In an attempt to make the calculation of contacts more manageable, in 2010 we built the SimiCon Web server.^12^ SimiCon (see Figure 1) is a free tool that allows for automatic calculation of equivalent contacts between two protein-ligand complexes and takes into account ligand and receptor symmetry (http://ablab.ucsd.edu/SimiCon). Thanks to SimiCon and other initiatives, interaction-based measures are progressively being incorporated in worldwide docking assessments.^13,14^

**Figure 1.**
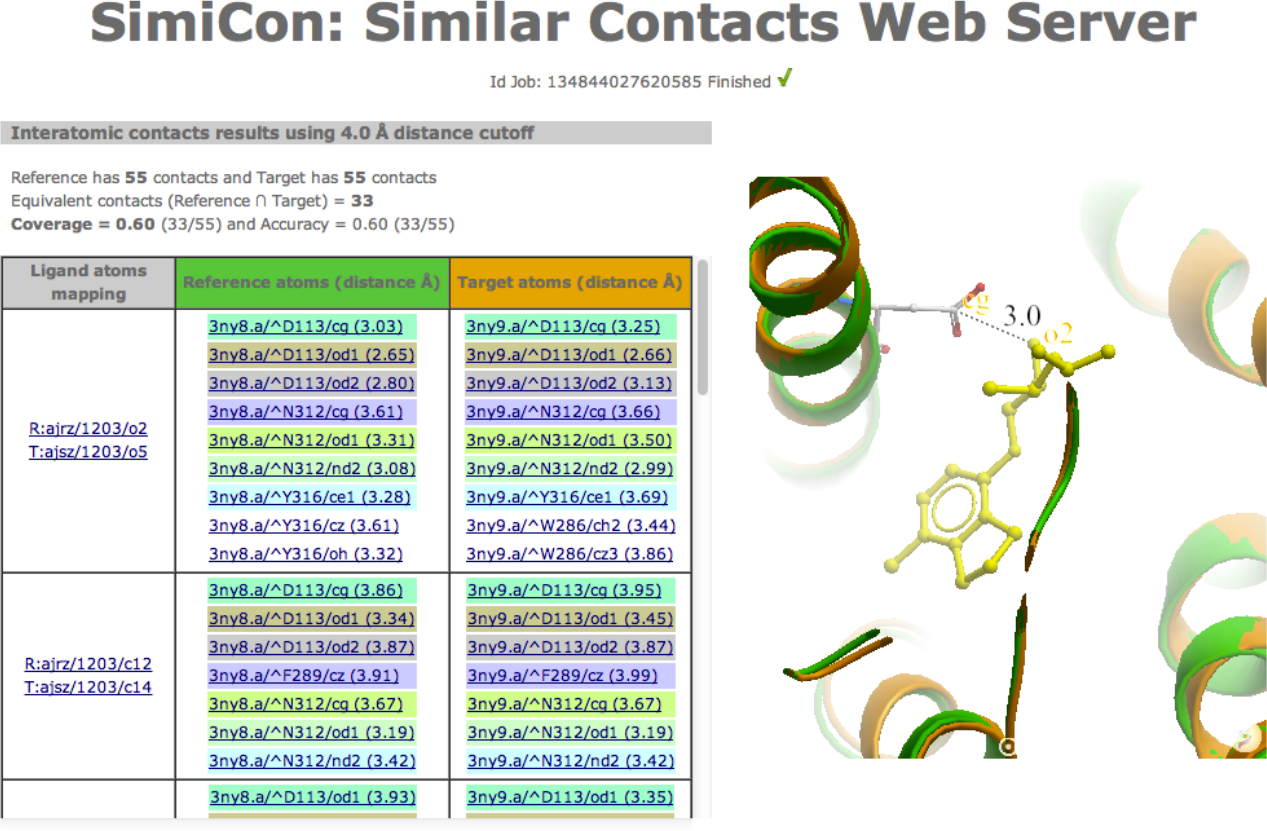
Screenshot showing results obtained with the SimiCon (Similar Contacts) web server (http://ablab.ucsd.edu/SimiCon). The server allows for calculation of equivalent contacts between two proteinligand complexes.

### 2.2 Ligand Screening

From the point of view of drug discovery, the real value of molecular docking relies on its capacity to filter *new* truly active molecules from inactive ones in massive VS experiments.^15^ Since most known ligands will never be crystallized,^16^ the screening capacity is benchmarked mostly by geometry-free descriptors. The assumption is that, providing that the docking tool has been parameterized for capturing key receptor-ligand interactions, the scores will be representative of the “correctness” of the poses. A widely used metric to quantify screening performance is the area under the receiver operating characteristic curve (ROC), abbreviated as AUC17 and its variations. The AUC plots the ratio of true positives (TP, *y*-axis) against the ratio of false positives (FP, *x*-axis). A perfect recognition will get an AUC value of 1.0 (100%), whereas a random selection will display an AUC of 0.5 (50%). Recently, we introduced the Normalized *SQ*uare root AUC (NSQ_AUC) metric, which as the log-AUC, BEDROC, RIE, SLR and the pROC meausres,^15, 18, 19^ puts emphasis on “early” hit enrichment.20 The NSQ_AUC measure returns a value of 1.0 for a perfect separation of signal from noise and values close to zero for a random selection

## 3 Single Receptor Conformation Docking

### 3.1 Self-docking

The most basic exercise for computing both the accuracy and the sampling thoroughness of any docking tool is to re-dock a compound into its cognate receptor. ICM was recently benchmarked for self-docking accuracy and screening as a part of a programmatic theme “Docking and Scoring: A Review of Docking Programs”, using the Astex,^21^ DUD^1^ and WOMBAT^22^ data sets. The developments and future prospects from such a campaign were presented during a symposium at the 241st ACS National Meeting held in March 2011 in Anaheim, CA, USA. With the 85 X-ray structures in the Astex data set, ICM performance in geometry prediction was satisfactory, placing near native poses at the top #1 in 91% of the cases and the top #3 in 95% of the cases. The remaining cases included problematic entries displaying water-mediated hydrogen bonds, metal coordination, or highly symmetrical or flexible compounds.^14^

### 3.2 Cross-Docking

Self-docking is a great exercise for calibration and benchmarking of the docking tools, but in practice all docking experiments are cross-docking ones. As discussed before, cross-docking validation is usually separated into two overlapping categories: one taking into account geometry predictions, and the other accounting for discrimination of binders from non-binders.

Prospective validation of docking poses with new crystallographic data, though existing ^23^, is rare. To this date, most geometry validations are performed with *existing* X-ray structures. In this sense, a key issue is the availability of curated data on which to perform the benchmarks. In our laboratory we have been compiling the Pocketome, an encyclopedia of conformational ensembles of all druggable binding sites that can be identified experimentally from co-crystal structures in the PDB (http://pocketome.org).^2^ The cross-docking performance of individual X-ray structures has been estimated at ~50%, or in other words, near native poses are obtained for ~ 50% of ligands when a single X-ray receptor is used.^24-27^

ICM virtual screening performance was also tested in the same 2011 exercise against the 40-target DUD and 11-target WOMBAT sets. VS using single pockets of the un-modified crystals provided a mean AUC value of 0.72 ± 0.16, with a minimum of 0.27 and a maximum of 0.96. The inclusion of receptor flexibility by a simple MC pocket refinement improved the early enrichments and overall discrimination, still using a single pocket per target (AUC values; mean: 0.79 ± 0.14, maximum: 0.98, minimum: 0.37).^14^

### 3.3 Screening performance variability among different X-ray structures of the same protein

A common phenomenon observed when working with Pocketome ensembles is that the docking and screening performance of individual pockets varies widely.^28^ In that regard, visual inspection, chemical intuition, or other descriptors (such as crystal resolution) are not enough to predict a given pocket's performance. An example of this effect can be observed in a recent study conducted with the Pocketome entry *ESR1_HUMAN_300_551* for the human estrogen receptor alpha (ERα), which by the time of this analysis consisted of 31 holo-structures. For each individual receptor, we performed retrospective flexible-ligand ICM-VS using a small ligand set consisting of the 31 cognate ligands as actives and 31 decoys (selected from the DUD entry for ERα). In this case we found that the best pocket displayed an NSQ_AUC = 0.80, and the worst an NSQ_AUC = 0.53, having an average value of 0.62 (see Figure 2).

**Figure 2.**
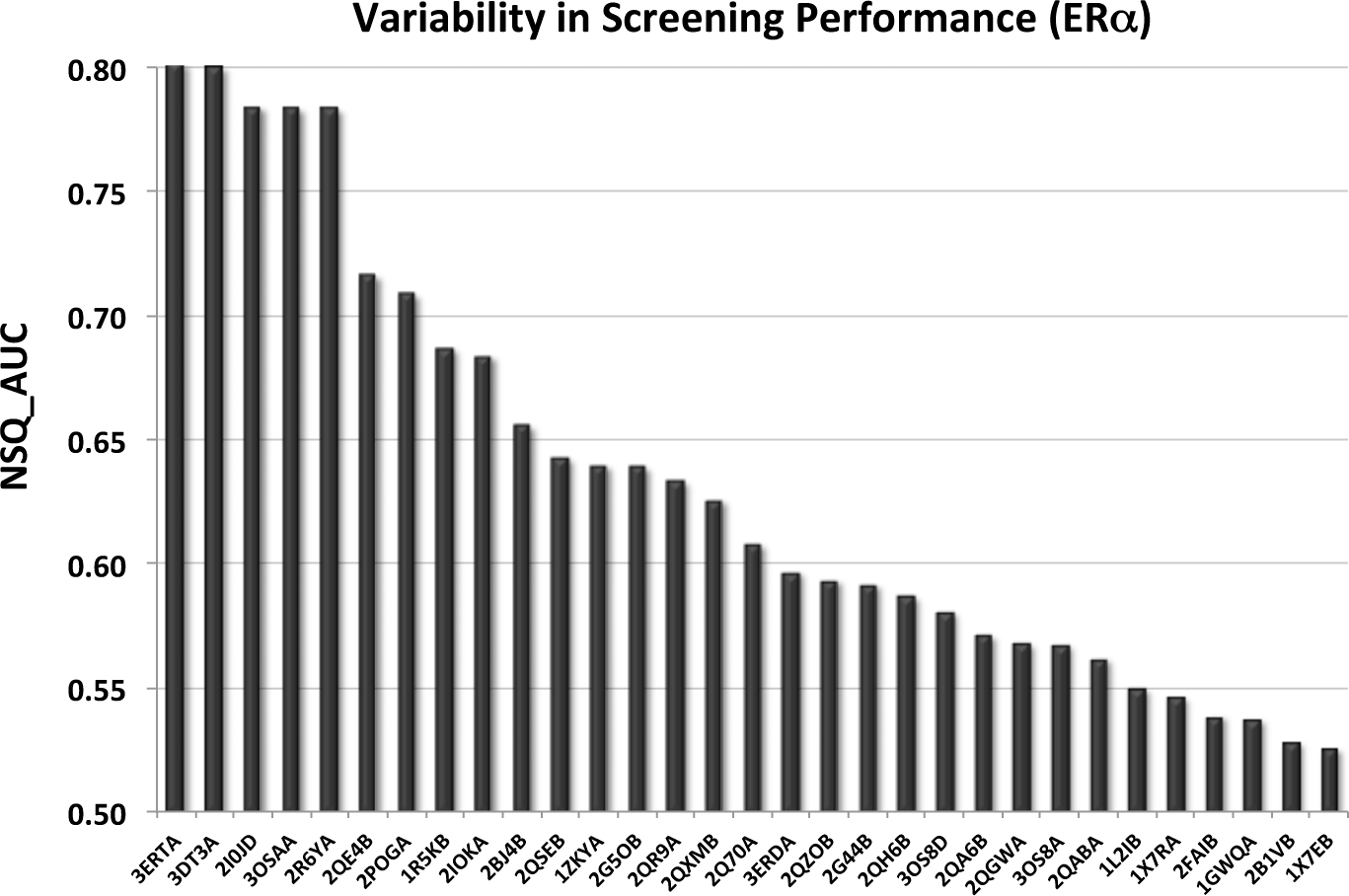
Variability in VS performance of the 31 PDB IDs from the *ESR1_HUMAN_300_551* Pocketome entry with a small ligand set (31 actives + 31 decoys). The performance was measured by the Normalized SQuare root Area Under the ROC Curve (NSQ_AUC).

### 3.4 Selecting the best individual pockets

As discussed above, predicting individual pocket performance without performing any docking is challenging. In practice, screening campaigns with small ligand sets are used to rank pockets according to their individual performance. For instance, with the *ESR1_HUMAN_300_551* Pocketome entry, the two best individual pockets (according to VS with a small test set of 31 actives + 31 decoys) 3ERT_A and 3DT3_A were also those performing better individually in a much larger set of unrelated compounds (16 true actives and 5586 inactives from ChEMBL).^29^ Unluckily, if activity data are scarce (or nonexistent) there are only a few guidelines for selecting the pockets (see Figure 3), like: to prefer holo-structures to apo ones, use only crystals with trusted resolution, or discard pockets consistently creating unfavorable scores (or strange ligand poses).^25, 30^

## 4 Multiple Receptor Conformations Docking: Rise and Fall

Simultaneous sampling of the protein and the ligand in docking is unreachable for screening purposes. As a practical alternative, ensemble docking has attracted the attention of many research groups because of its speed and its straightforward implementation. In ensemble docking, the receptor plasticity is introduced by using multiple discrete conformers that can have experimental or in silico origin.^31^ Each receptor will undergo an independent docking run, usually performed in multi-CPU desktop computers or Linux clusters. After the docking, the scores (or ranks) from individual runs are combined and only the best values are kept to compute the AUC (or derivatives of the AUC).^30^

Ensemble docking still may represent a hurdle for laboratories not disposing enough computational resources. To alleviate that situation, we recently developed the four-dimensional docking (4D-docking), a method that reduces the sampling time while preserving the accuracy of traditional ensemble docking ^27^. The foundation of the method is to add the maps of each receptor as an extra dimension to the MC search space, allowing the entire ligand to move from one 4D “plane” to another, thus switching between different receptor conformations. In a recent comparison of our three cross-docking approaches: single receptor docking, multiple receptor conformation docking and 4D docking in a compiled benchmark (100 proteins, >1000 structures) we obtained success rate of 46.6%, 79.6%, and 77.3%, respectively. Apart from our own findings, the literature contains many successful examples on the advantages of ensemble docking with respect to single receptor conformation docking with experimental and in silico conformers (we suggest reading the reference^31^). Apart from improving the separation of actives from inactives, the use of ensembles helps capture a wider scaffold diversity,^30^ and helps profile compounds according to affinity.^32,33^

Despite the euphoria, ensemble docking has not reached the desired potential because too many pockets increase the ratio of false positives, thus deteriorating the performance.^25, 28^

### 4.1 Selecting a “team” of complementary pockets

Those opting for the ensemble docking route will be confronted with the issue of optimal pocket selection. To address this hurdle, a key point is the availability of activity data for several compounds (see Figure 3).

Luckily, for targets with no data available for active ligands (often these cases are the most appealing from a pharmaceutical point of view), ensemble docking seems a reasonable choice. Several studies have demonstrated that ensembles of randomly chosen pocket structures on average outperform individual ones.^25, 28, 34^ Intriguingly, the ensemble tolerance to the addition of pockets is finite, being ~3-5 the “magic” number of structures acting synergistically.^25^ For targets where information on a handful of active ligands exists (ideally co-crystal structures), a few extra guidelines can help us narrow the selection to the essential conformers. For instance, we recommend selecting holo-structures and discarding apo ones, discarding pockets consistently creating unfavorable scores (or strange ligand poses), selecting those pockets with the largest ligands or cavities (in principle they will accommodate large *and* small compounds), pocket clustering, or selecting pockets according to ligand similarity.^35, 36^ Recently it has been suggested that optimal ensemble selection protocols are scoring function dependent.^34^

**Figure 3.**
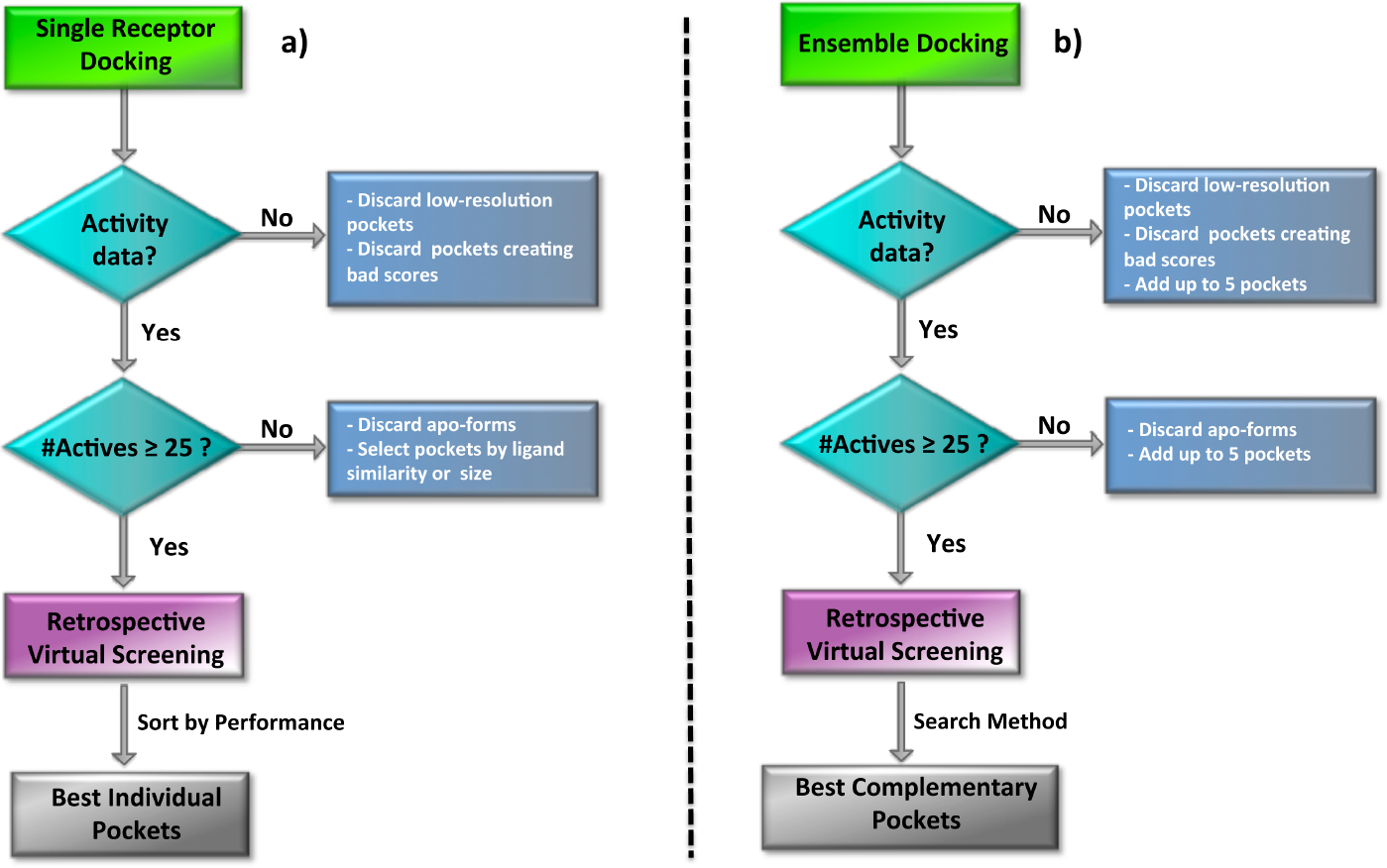
Scheme summarizing the strategies to follow for selecting the best individual pockets (a) or the best ensemble (b) for VS.

The best possible scenario appears when ~ 25 or more actives are known, especially if the compounds displayed decent potency in experimental studies. As with individual pocket selection, we recommend performing retrospective campaigns with known actives plus decoys (at different ratios, if possible). Unfortunately, ensembles created by the above rules, or created by the addition of the best *individual* pockets do not necessarily lead to a better ensemble performance.^25^ Thus, to find the optimal combination of pockets we should rely on some sort of “brute force” search method. In that regard, an obvious choice is to perform an exhaustive combinatorial search.^28, 34^ This approach, however, is only computationally feasible for relatively small ensembles, becoming prohibitive for large ones. To address the speed of the calculation, alternative methods have been developed.^37^ Very recently, we proposed a very efficient selection strategy, which performs on par with the exhaustive search but in a fraction of the time. The method uses a dual strategy, where an exhaustive search is performed until *M* = 2 *(M* is the number of complementary pockets) plus individual pocket addition when *M* ≥ 3. The search procedure is repeated until the addition of pockets no longer increases the value of the fitness function, or until *M* reaches the value set by the user.^29^ We tested our selection method with the 31 X-ray structures in *ESR1_HUMAN_300_551* Pocketome entry and found that an ensemble consisting of 7 pockets provided the maximum discrimination as measured by the NSQ_AUC (see Figure 4). Not all the 8 pockets contributed equally to the recognition, and only 5 added significant value to the separation of actives from decoys.

**Figure 4.**
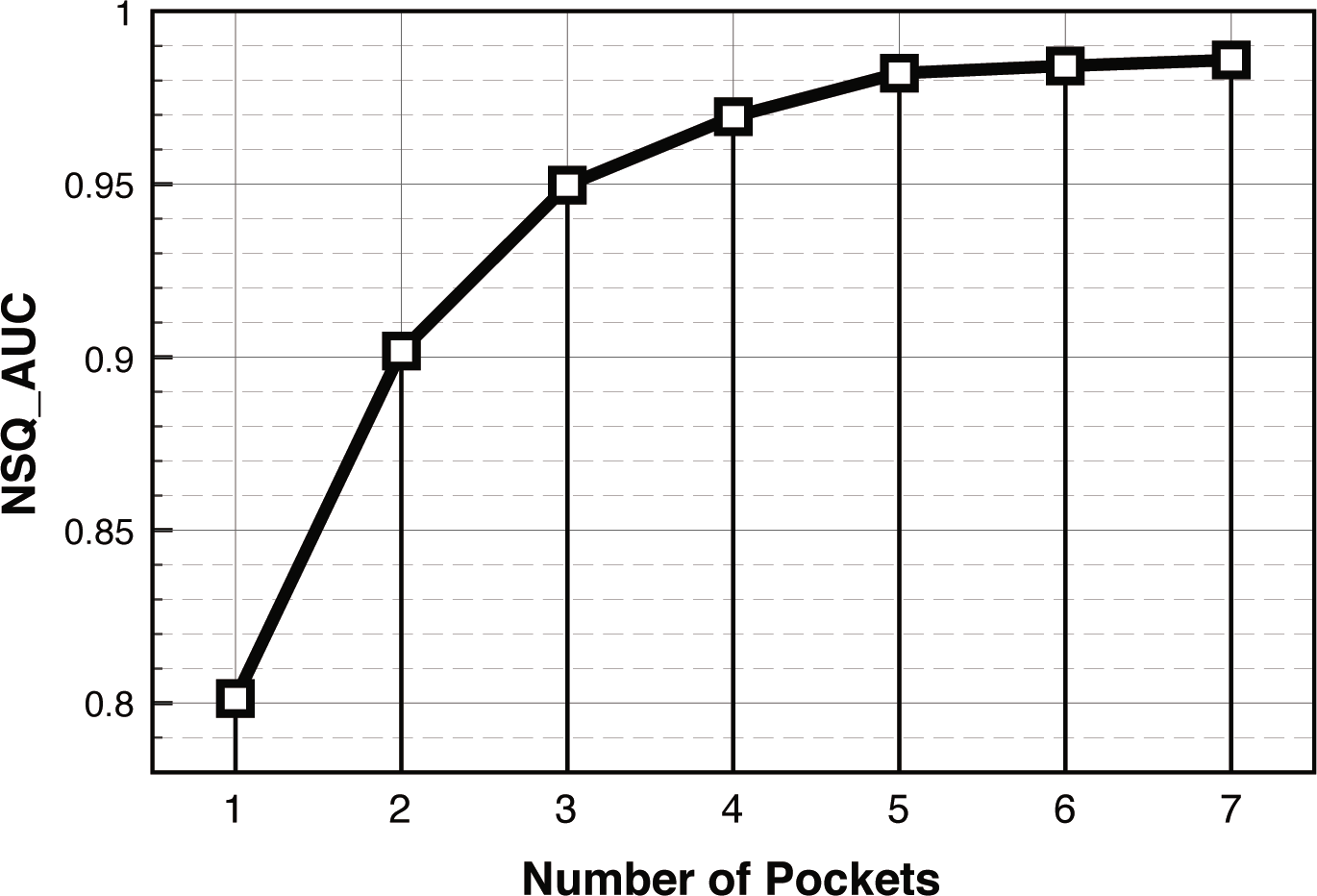
Increase of NSQ_AUC values by the addition of complementary pockets. The results correspond to the “team” selection method used with the 31 X-ray structures from the *ESR1_HUMAN_300_551* Pocketome entry. The ligand set consisted of the 31 cognate ligands as actives plus 31 decoys from the DUD.

## 5 Alibero: Improving Docking Performance of X-Ray Structures and Homology Models

In the context of ensemble docking, we recently published ALiBERO (Automatic Ligand-guided Backbone Ensemble Receptor Optimization), a new computational tool created to improve the discrimination power of X-ray structures and homology models. Starting from single or multiple receptor structures, the software iteratively creates receptor ensembles, performs VS docking and selects the “team” of complementary pockets that maximizes the recognition of ligand actives from decoys (see Figure 5). The iterative sampling-selection is able to improve the discriminative power of the models due to inheritance of conformational features, combined with newly found features.

**Figure 5.**
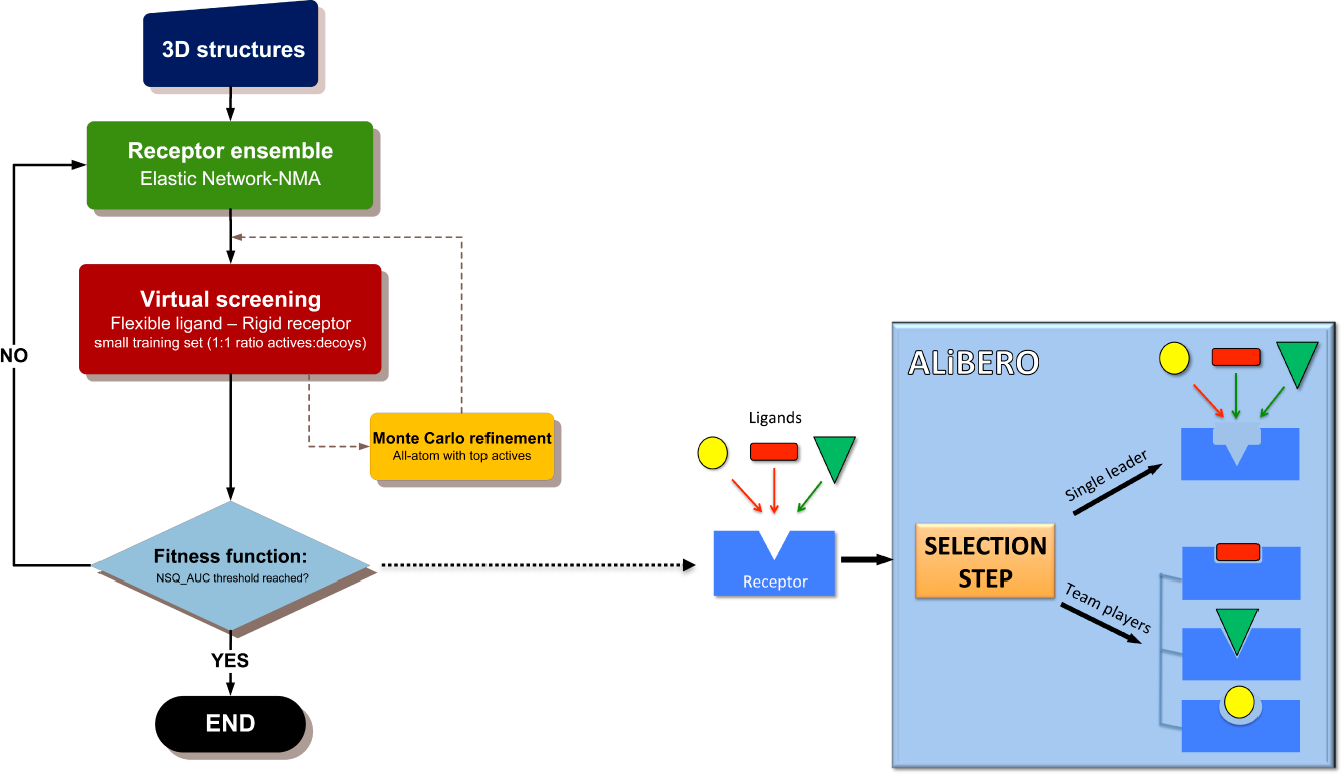
Flowchart representing ALiBERO algorithm and the two strategies used for the selection of “children” pockets. In the “single leader” mode only the best individual pocket will be selected for the next generation, whereas in the “team players” mode the best ensemble will be selected. As input, the algorithm takes one or multiple 3D receptor structures plus a ligand training set consisting of target-specific ligands and decoys.

ALiBERO is currently being tested in multiple Pocketome entries including popular pharmaceutical targets (e.g., nuclear receptors, kinases, phospathases, GPCRs) and in distant homology models of trans membrane proteins yielding encouraging results. The software is rapidly becoming a routine tool within the ICM-docking community, both in academia and industry. The script is available from the authors upon request as a free add-on to ICM (Molsoft LLC) molecular modeling package for the Linux platform.

## Acknowledgments

The authors thank Karie Wright for help with manuscript preparation. MR is supported by a Marie Curie OIF fellowship. This work was supported by NIH grant 1-R01-GM074832.

